## Supplementary files for "Distinct representational properties of cues and contexts shape fear and reversal learning"

**Supplementary Table 1.** Wilcoxon post-hoc tests, with W and corrected p values, for the pairwise comparisons of US expectancy between each CS type for each learning phase.

| Phase | Comparison | W | p |
|---|---|---|---|
| **Acquisition** | CS++ vs CS+- | W=190.0 | p=0.39 |
| **Acquisition** | CS++ vs CS-+ | W=26.5 | p=1.99e-06 |
| **Acquisition** | CS++ vs CS-- | W=8.0 | p=4.65e-08 |
| **Acquisition** | CS-+ vs CS+- | W=31.0 | p=4.42e-06 |
| **Acquisition** | CS-+ vs CS-- | W=150.5 | p=0.09 |
| **Acquisition** | CS+- vs CS-- | W=3.0 | p=9.31e-09 |
| **Reversal** | CS++ vs CS+- | W=96.5 | p=0.009 |
| **Reversal** | CS++ vs CS-+ | W=106.0 | p=0.01 |
| **Reversal** | CS++ vs CS-- | W=18.5 | p=9.42e-07 |
| **Reversal** | CS-+ vs CS+- | W=137.0 | p=0.08 |
| **Reversal** | CS-+ vs CS-- | W=18.5 | p=1.75e-05 |
| **Reversal** | CS+- vs CS-- | W=33.5 | p=1.19e-05 |
| **Test new** | CS++ vs CS+- | W=52.5 | p=0.001 |
| **Test new** | CS++ vs CS-+ | W=100.0 | p=0.001 |
| **Test new** | CS++ vs CS-- | W=19.5 | p=1.94e-05 |
| **Test new** | CS-+ vs CS+- | W=202.5 | p=0.75 |
| **Test new** | CS-+ vs CS-- | W=28.0 | p=4.32e-05 |
| **Test new** | CS+- vs CS-- | W=46.5 | p=0.0002 |
| **Test old** | CS++ vs CS+- | W=95.5 | p=0.008 |
| **Test old** | CS++ vs CS-+ | W=161.5 | p=0.22 |
| **Test old** | CS++ vs CS-- | W=48.0 | p=0.0002 |
| **Test old** | CS-+ vs CS+- | W=158.0 | p=0.2 |
| **Test old** | CS-+ vs CS-- | W=21.0 | p=2.19e-05 |
| **Test old** | CS+- vs CS-- | W=71.5 | p=0.001 |

**Supplementary Table 2.** Cluster statistics for the cue generalization (CS+ > CS−) contrast in fear acquisition. FWER-corrected p-values of the clusters are marked as significant according to a Bonferroni-corrected threshold of **$p < 0.025$ (0.05/2)**.

| Contrast | Cluster ID | Peak x | Peak y | Peak z | Mean cluster t-value | p-value | Significant (Bonferroni) | Volume (mm³) | Brain region |
|---|---|---|---|---|---|---|---|---|---|
| **CS+ > CS−** | 1 | 3.5 | 25 | 56.5 | 4.38 | 0.00022 | Yes | 4515.63 | Superior Medial Frontal (Left) |
| | 2 | -19 | -82.5 | -28.5 | 4.33 | 0.00025 | Yes | 3937.50 | Cerebellum Crus I (Left) |
| | 3 | 28.5 | -70 | -51 | 4.43 | 0.00019 | Yes | 3625.00 | Cerebellum Lobule VIII (Right) |
| | 4 | 56 | 5 | 39 | 4.37 | 0.00022 | Yes | 3156.25 | Inferior Frontal Operculum (Right) |
| | 5 | 58.5 | 5 | -26 | 4.41 | 0.00020 | Yes | 2484.38 | Middle Temporal Gyrus (Right) |
| | 6 | -61.5 | -15 | 41.5 | 4.28 | 0.00028 | Yes | 2281.25 | Postcentral Gyrus (Left) |
| | 7 | 63.5 | -25 | 26.5 | 4.41 | 0.00020 | Yes | 1984.38 | Supramarginal Gyrus (Right) |
| | 8 | -31.5 | 20 | -6 | 4.26 | 0.00030 | Yes | 1656.25 | Insula (Left) |
| | 9 | -26.5 | -45 | -53.5 | 4.63 | 0.00012 | Yes | 1562.50 | Cerebellum Lobule VIII (Left) |
| | 10 | -39 | 12.5 | 31.5 | 4.23 | 0.00032 | Yes | 1531.25 | Inferior Frontal Operculum (Left) |
| | 11 | 56 | -47. | 19 | 4.13 | 0.00041 | Yes | 1453.13 | Superior Temporal |

| | | | | | | | | | |
|---|---|---|---|---|---|---|---|---|---|
| | | | 5 | | | | | | Gyrus (Right) |
| | 12 | 43.5 | -7.5 | 59 | 4.25 | 0.00030 | Yes | 1312.50 | Precentral Gyrus (Right) |
| | 13 | -21.5 | -65 | 36.5 | 4.09 | 0.00045 | Yes | 1109.38 | Superior Parietal Lobule (Left) |
| | 14 | 41 | 17.5 | -11 | 4.25 | 0.00030 | Yes | 953.13 | Insula (Right) |
| | 15 | -56.5 | 10 | -21 | 4.28 | 0.00028 | Yes | 921.88 | Middle Temporal Gyrus (Left) |
| | 16 | 3.5 | -27.5 | 66.5 | 4.26 | 0.00030 | Yes | 906.25 | Paracentral Lobule (Right) |
| | 17 | -11.5 | 5 | 19 | 4.38 | 0.00022 | Yes | 812.50 | Caudate (Left) |
| | 18 | -44 | -55 | 54 | 4.29 | 0.00027 | Yes | 750.00 | Inferior Parietal Lobule (Left) |

**Supplementary Table 3.** Cluster statistics for cue generalization and item stability in fear reversal. FWER-corrected p-values of the clusters are marked as significant according to a Bonferroni-corrected threshold of **$p < 0.00625$ (0.05/8)**.

| Contrast | Cluster ID | Peak x | Peak y | Peak z | Mean clust | p-value | Significant (Bonferro | Volume (mm³) | Brain region |
|---|---|---|---|---|---|---|---|---|---|

| | | | | | er t-value | | ni) | | |
|---|---|---|---|---|---|---|---|---|---|
| **CS++ > CS− cue generalization** | 1 | 6 | 50 | 39 | 4.19 | 0.00039 | Yes | 906.25 | Superior Medial Frontal (Right) |
| **Current CS+ > CS− cue generalization** | 1 | 38.5 | 22.5 | -8.5 | 4.08 | 0.00046 | Yes | 1375.00 | Inferior Frontal Triangular (Right) |
| | 2 | -41.5 | 57.5 | 9 | 4.49 | 0.00015 | Yes | 1109.38 | Middle Frontal Gyrus (Left) |
| | 3 | 3.5 | -70 | 39 | 3.98 | 0.00055 | Yes | 734.38 | Precuneus (Right) |
| | 4 | -34 | -62.5 | 54 | 4.20 | 0.00038 | Yes | 718.75 | Superior Parietal Lobule (Left) |
| **CS-changing vs CS-consistent item stability** | 1 | -1.5 | -75 | 31.5 | 4.34 | 0.00018 | Yes | 3796.88 | Precuneus (Left) |
| | 2 | 46 | 40 | -6 | 4.35 | 0.00017 | Yes | 671.88 | Inferior Frontal Orbital (Right) |

**Supplementary Table 4.** Cluster statistics for item stability in test new and test old. FWER-corrected p-values of the clusters are marked as significant according to a Bonferroni-corrected threshold of **p < 0.00625 (0.05/8)**.

| Contrast | Cluster ID | Peak x | Peak y | Peak z | Mean cluster t-value | p-value | Significant (Bonferroni) | Volume (mm³) | Brain region |
|---|---|---|---|---|---|---|---|---|---|

| | | | | | | | | | |
|---|---|---|---|---|---|---|---|---|---|
| **Test new: CS+ → CS++** | 1 | -64 | -12.5 | -13.5 | 4.29 | 0.00023 | Yes | 812.50 | Middle Temporal (Left) |
| **Test old: CS++ > CS−−** | 1 | -26.5 | -27.5 | -28.5 | 4.52 | 0.00013 | Yes | 1062.50 | Fusiform (Left) |

**Supplementary Table 5.** Cluster statistics for context specificity differences between acquisition and reversal. P-values are based on a two-tailed paired t-test with 24 participants. Clusters are marked as significant according to a Bonferroni-corrected threshold of **$p < 0.00714$ (0.05/7)**.

| Cluster ID | Peak x | Peak y | Peak z | Mean cluster | p-value | Significant (Bonferroni) | Volume (mm³) | Brain region |
|---|---|---|---|---|---|---|---|---|

| | | | | t-value | | | | |
|---|---|---|---|---|---|---|---|---|
| **1** | 3.5 | 60 | 16.5 | -4.56 | 0.00012 | Yes | 5031.25 | Superior Medial Frontal (Right) |
